# A gene-level methylome-wide association analysis identifies novel Alzheimer’s disease genes

**DOI:** 10.1101/2020.07.13.201376

**Authors:** Chong Wu, Jonathan Bradley, Yanming Li, Lang Wu, Hong-Wen Deng

## Abstract

**Motivation:** Transcriptome-wide association studies (TWAS) have successfully facilitated the discovery of novel genetic risk loci for many complex traits, including late-onset Alzheimer’s disease (AD). However, most existing TWAS methods rely only on gene expression and ignore epigenetic modification (i.e., DNA methylation) and functional regulatory information (i.e., enhancer-promoter interactions), both of which contribute significantly to the genetic basis of AD.

**Results:** This motivates us to develop a novel gene-level association testing method that integrates genetically regulated DNA methylation and enhancer-target gene pairs with genome-wide association study (GWAS) summary results. Through simulations, we show that our approach, referred to as the CMO (cross methylome omnibus) test, yielded well controlled type I error rates and achieved much higher statistical power than competing methods under a wide range of scenarios. Furthermore, compared with TWAS, CMO identified an average of 124% more associations when analyzing several brain imaging-related GWAS results. By analyzing to date the largest AD GWAS of 71,880 cases and 383,378 controls, CMO identified six novel loci for AD, which have been ignored by competing methods.

**Availability and implementation:** Software: https://github.com/ChongWuLab/CMO

**Supplementary information:** Supplementary data are available at Bioinformatics online.

## Introduction

Alzheimer’s disease (AD) is a progressive loss of memory and cognition, for which no effective treatment or cure currently exists [1]. The main hallmarks of AD are amyloid plaques and neurofibrillary tangles, which might develop 20 years or more before the onset of rapid neurodegeneration and cognitive symptoms [1]. Thus, effective and early assessment of AD risk is critical to identify individuals at high risk and decrease the public health burden of AD. While A*β* and tau levels in cerebrospinal fluid (CSF) are considered to be potential markers for assessing AD risk, lumbar puncture is painful and routine check of CSF is difficult [2]. On the other hand, several studies [3, 4] have suggested specific genes in blood can serve as candidate biomarkers for risk assessment. Thus, a better understanding of AD-associated genes in blood, beyond the scope of current understanding, may lead to a more powerful risk assessment.

One strategy to identify associated genes is to use gene-level integrative association tests [5–11], which aggregate potential regulatory effects of individual genetic variants across a gene of interest to test gene-trait associations. One such family of tests is transcriptome-wide association studies (TWAS) [5–7, 10], which integrate expression reference panels (i.e., expression quantitative trait loci [eQTL] cohorts with matched individual-level expression and genotype data) with genome-wide association studies (GWAS) data to discover gene-trait associations. TWAS have garnered substantial interest within the human genetics community, have identified many novel trait-associated genes, and have deepened our understanding of genetic regulation in many complex traits, including AD [12].

There is an exciting opportunity to develop novel and powerful integrative association tests by incorporating additional functional and epigenetic information. Specifically, there has been an increasing interest in the role of epigenetic modifications (such as DNA methylation [DNAm]) that interact between genome and environment in complex disease etiology [13, 14]. DNAm is essential for mammalian development and mediates many critical cellular processes, including gene regulation, genomic imprinting, and X chromosome inactivation [15]. Recent epigenome-wide association studies (EWAS) [16] have led to a rapid increase in identifying robust associations between DNAm and complex traits, including AD [17]. Critically, many of the identified significant CpG sites (CpGs) are different from the well-established AD genetic risk loci identified by GWAS. This highlights the potential utility of EWAS in characterizing additional insights into AD pathogenesis and risk assessment [18]. However, the causal relationship between DNAm and AD is difficult to establish because the DNAm levels are affected by environmental and lifestyle factors [19]. To reduce limitations such as reverse causation and unmeasured confounders that are commonly encountered in conventional EWAS, methylome-wide analyses (MWAS) [20, 21] have been proposed to test associations between genetically regulated DNAm and the trait of interest. However, existing methods are largely focused on individual CpG by testing each CpG separately. Additionally, we and others have shown that integrating enhancer-promoter interactions will lead to improved statistical power for gene-level association tests [22] as enhancers are key gene-regulatory DNA sequences that control gene expression by engaging in physical contacts with their cognate genes [23]. However, it is still unclear how to effectively integrate genetically regulated DNAm and promoter-enhancer interactions with GWAS results.

In this work, we propose a novel computationally efficient gene-level association testing method, termed cross-methylome omnibus (CMO) test. CMO integrates genetically regulated DNAm in enhancers, promoters, and the gene body to identify additional disease associated genes that are ignored by existing methods. In an applied analysis for AD risk, for generating candidates that may facilitate risk assessment for AD, we focused on a set of prediction models for blood tissue. Through simulations and analysis of several brain imaging-derived phenotypes, we demonstrate that CMO achieves high statistical power while controlling the type I error rate well. Additionally, by reanalyzing several AD GWAS summary datasets [24–26], we demonstrate that our approach can reproducibly identify additional AD-associated genes that are not able to be identified by the competing methods. These newly identified genes in blood may serve as promising candidates for risk assessment of AD.

## Materials and methods

### Overview of CMO test

We build upon previous work [5–7, 11, 20, 21] to develop a novel integrative gene-based test for identifying novel genes that may influence the trait of interest through DNAm pathways. CMO involves three main steps. First, CMO links CpGs located in enhancers, promoters, and the gene body (including both exons and introns) to a target gene because DNAm in enhancers and promoters may play vital roles in gene regulation [27]. Of note, CMO integrates and thus utilizes much more comprehensive enhancer-promoter interaction information from a comprehensive database [28], which is different from our previous work [11, 22] that only integrates two sets of enhancer-promoter interaction data for illustration purposes. Second, by leveraging existing DNAm prediction models (built from reference datasets that have both genetic and DNAm data), CMO tests associations between genetically regulated DNAm of each linked CpG and the trait of interest using a weighted gene-based test. Because the underlying genetic structure is unknown, different CpG-based tests may be more powerful under different scenarios. To offer consistently high statistical power, we apply a Cauchy combination test [29] to combine the results from a set of widely used tests. Third, CMO applies a Cauchy combination test to combine statistical evidence from multiple CpGs for a target gene to test whether the target gene is associated with the trait of interest.

### Brief overview of Aggregated Cauchy Association Test (ACAT)

ACAT [29, 30] is a recently proposed Cauchy combination test to combine *p*-values. Because ACAT is an important building block for our proposed method CMO, we briefly review it here.

Suppose *p*_1_, *p*_2_,*…*, *p_k_* denote *k p*-values to be combined by ACAT. For testing the association between genetically regulated DNAm levels and the trait of interest, the *p* values correspond to *k* genetic variants with non-zero weights, which are either provided by GWAS summary results or can be calculated from a *Z* score. For combining results from multiple CpGs, *p_j_* can be the *p*-value of the *j*th CpG. ACAT first transforms the original *p*-values to standard Cauchy random variables under the null hypothesis and then uses the weighted summation of transformed *p*-values as the test statistic, where flexible non-negative weights are allowed. Specifically, ACAT uses the following test statistic

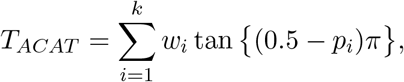

where the *p_i_*’s are the *p*-values and the *w_i_*’s are external non-negative weights. Under any arbitrary correlation structure among the *p*-values, *T_ACAT_* can be approximated using a Cauchy distribution and the *p*-value of ACAT can be approximated by [29, 30],

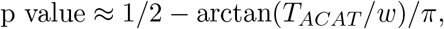

where 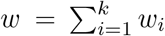. In other words, ACAT circumvents the challenging and computationally extensive step of estimating correlations among the *p*-values, which is often needed for competing methods.

### Details of CMO test

The aim of our method CMO is to identify associated genes that affect the trait of interest through DNA methylation pathways. We consider only one gene for illustration because the same procedure can be applied for each gene sequentially. The null hypothesis is that the gene to be tested is not associated with the trait of interest. CMO involves the following three steps.

#### Step 1: Linking CpG sites (CpGs) to a target gene

We are motivated to link CpGs that are located in the enhancers, promoters, and the gene body to their target genes. This is because CpGs in enhancers and promoters can play important roles in gene regulation [27]. Following our previous work [11], we define enhancers, promoters, and gene body as follows. A gene body region includes all the introns and exons of a target gene. To include cis-acting regulatory regions and alleviate the burden of determining gene direction, two promoters of a target gene are defined as a 500 bp extension [31] on either side of the gene body region beyond its transcription start site (TSS) and transcription end site (TES), respectively. For enhancers, we use an integrated database called *GeneHancer* [28]. By integrating reported enhancers from four genome-wide databases (the ENCODE, the Ensembl regulatory build, the FANTOM, and the VISTA Enhancer browser) and linking enhancers to their target genes via several complementary approaches, GeneHancer provides comprehensive enhancer-gene information [28].

The genomic coordinates of genes are obtained from the human genome assembly GRCh37/hg19. We use the Illumina HumanMethylation450 v1.2 manifest file to determine the position of CpGs and map/link CpGs in the gene body, two promoters, and enhancers regions to its target gene accordingly. We assume we map *q* CpGs to a target gene.

#### Step 2: Testing for each linked CpG

Similar to conventional TWAS framework, Baselmans et al. [21] built prediction models for predicting genetically regulated DNAm. Specifically, at a false discovery rate (FDR) of 5%, there were 151,729 CpGs with at least one significant DNAm quantitative trait locus (mQTL). For these 151,729 CpGs, a Lasso model was applied with local single nucleotide polymorphisms (SNPs) (within 250 Kb extension) were chosen as predictors, and DNA methylation level was defined to be the outcome. Baselmans et al. [21] further provided the mQTL-derived weights ****W**** = (*W*_1_, …, *W*_*k*_)^′^ and corresponding LD matrix ****V**** for local SNPs as a publicly available resource, on which CMO was based.

We focus on one CpG for illustration. Suppose we have the corresponding *Z* score vector ****Z**** = (*Z*_1_, …, *Z*_*k*_)^′^ for the CpGs being considered. Under the null hypothesis (no association), ****Z**** follows a normal distribution with mean 0, that is ***Z*** ∼ *N* (0, ***V***). To test the association between the estimated genetically regulated DNAm and the trait of interest, Baselmans et al. proposed the MWAS test statistic [21]

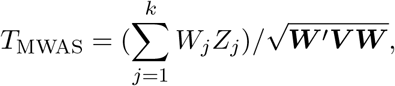

where *W*_*j*_ is the mQTL-derived weight for SNP *j*, provided by Baselmans et al. [21]. MWAS can be viewed as a standardized weighted burden test. The statistic (*T*_MWAS_) has an asymptotic standard normal distribution, and its *p*-value can be computed analytically.

By over-weighting the methylation-associated SNPs, MWAS improves the statistical power and reduces the concern of a reverse-causality problem [21]. While appealing, MWAS also has similar limitations as TWAS [7, 32]. Specifically, the burden test is derived under the over-simplifying assumption that all (weighted) SNPs have the same effect sizes. Thus, the burden test (i.e., MWAS) will lose power if the effect directions of the (weighted) SNPs are different or the effect sizes are sparse (i.e., with many 0’s). Other widely used tests, such as SSU [33] and ACAT [29, 30], will be powerful under many scenarios, since the underlying true model is unknown and no uniformly most powerful test exists[34].

As a result, besides applying the burden test, we apply the weighted SSU test and ACAT. The SSU test statistic is,

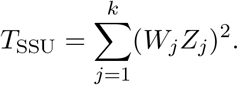

*T*_SSU_ follows a mixture of chi-square distribution asymptotically, and we can calculate its *p*-value analytically [33]. The ACAT test statistic is,

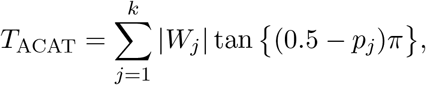

where *p*_*j*_ is the *p*-value for SNP *j*. We use the absolute value of *W*_*j*_ as the weight for SNP *j* since ACAT requires non-negative weights. As discussed previously, the *p*-value can be calculated analytically.

Because there is no uniformly most powerful test in general, and the optimal test depends on the underlying truth [34], which varies in practice, we apply the Cauchy combination test [29] to offer robust statistical power. Specifically, we define the test statistics as,

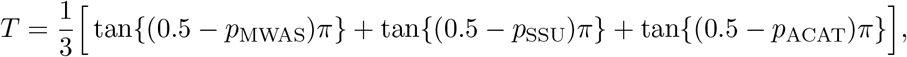

where *p*_MWAS_, *p*_SSU_, and *p*_ACAT_ are the *p*-values for the MWAS, SSU, and ACAT tests, respectively. Then the corresponding *p*-value is calculated as 0.5−{arctan(*T*)}/*π*. We repeat the above procedure for each linked CpGs of a target gene and obtain *P* = (*p*_1_*, …, p_q_*)′ for the *q* CpGs defined in Step 1.

#### Step 3: Integrating over multiple CpGs in a target gene

To robustly aggregate information from different CpGs, we propose an omnnibus test called cross-methylome omnibus (CMO) tests that applies the Cauchy combination test [29, 30] to the linked CpGs. The test statistics is

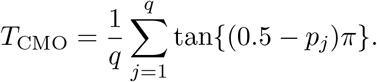

*T*_CMO_ can be approximated by a standard Cauchy distribution well and the *p*-value can be calculated as 0.5 − {arctan(*T*)}/*π* [29]. Of note, the approximation error goes to zero as the *p*-value of CMO goes to zero [29, 30]. In other words, the *p*-value calculation is particularly accurate when CMO has a highly small *p*-value, which is a useful feature in gene-based analyses because only highly small *p*-values can pass the stringent Bonferroni significance threshold and are of interest.

### Simulation settings

We conducted simulations to compare the performance of CMO and competing methods regarding the type I error rates and statistical power. Specifically, we used data from UK Biobank (application number 48240) and randomly chose 10,000 independent White British individuals. The imputed genetic data for 5,911 cis-SNPs (with minor allele frequency (MAF) > 1%, Hardy-Weinberg *p*-value > 10^−6^, and imputation “info” score > 0.4) of the most significant AD-associated gene *APOE* were used in the simulation studies.

We first simulated both genetically regulated DNAm values at different CpGs and genetically regulated gene expressions at different tissues. We randomly selected *m* CpGs and *m* tissues. Then the cis component of DNAm values in CpG *i* and cis component of gene expression in tissue *j* was generated as 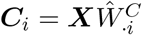 and 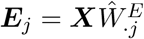, respectively. ****X**** is the *n × p* centered and standardized genotype matrix, and we used real effect size *Ŵ* to preserve the genetic architecture. Next, we simulated the quantitative trait by 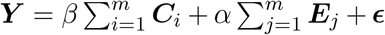, where 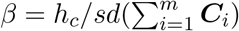 is the effect of CpGs on the trait, 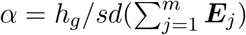 is the effect of gene expressions on the trait, ***ϵ*** ∼ *N* (0*, σ*^2^) is random noise, and *σ*^2^ determines the heritability of all the CpGs and gene expression in different tissues being considered. *h*_*c*_ and *h*_*g*_ controls the relative heritability contribution and effect size direction of CpGs and gene expressions, respectively. We performed an association scan on simulated data (***Y***, ***X***) and computed *Z* scores by a linear regression.

To evaluate type I error, we set *h*_*c*_ = 0 and *h*_*g*_ = 0, i.e., no association between the gene and trait. To evaluate power, we considered several combinations of *h*_*c*_ and *h*_*g*_. For example, we set *h*_*c*_ = −1 and *h*_*g*_ = 1 as methylation usually represses gene expression. We also considered diverse disease architectures by varying heritability (i.e., 0.005 and 0.01). We repeated simulations 500,000 times under the null and 1,000 times under the alternatives, and compared our proposed CMO test with standard TWAS and a multi-tissue TWAS, called MultiXcan [35]. Furthermore, we also considered taking the union of CpGs in gene body and promoter regions while applying an additional Bonferroni correction, denoted as union of MWAS or MWAS for simplicity. Statistical power was calculated as the proportion of 1,000 repeated simulations with *p*-value reaching the genome-wide significance threshold 0.05/20, 000 = 2.5 *×* 10^−6^.

### Application to AD GWAS data

To illustrate the proposed methods and deepen our understanding of genetic regulation in AD, we applied different gene-based association tests to identify AD risk genes by reanalyzing three sets of AD GWAS summary datasets: a meta-analyzed AD GWAS dataset with 17,008 AD cases and 37,154 controls [24], denoted as IGAP1; a genome-wide association study by proxy (GWAX) results with 14,482 (proxy) AD cases and 100,082 (proxy) controls the UK Biobank [25], denoted as GWAX; and the largest available AD GWAS data of 71,880 (proxy) AD cases and 383,378 (proxy) controls of European ancestry [26]. Of note, the proxy AD cases were defined by family history and showed quite strong genetic correlation with AD (*r*_*g*_ = 0.81) [26].

We performed a multi-stage gene-level association study for AD. In the performance evaluation stage, we identified significantly associated genes in the IGAP1 data set. Next, we replicated our findings by independent data from GWAX and then further validated by GWAS Catalog [36]. The *P*-values for the replication rate were calculated by a hypergeometric test, with the background probabilities estimated from all the genes being tested. In the end, we applied CMO and competing methods to the largest available AD GWAS dataset [26] for new discovery. We further performed the “Core Analysis” in Ingenuity Pathway Analysis (IPA) [37] for the identified associated genes to assess the enriched pathways, diseases, and networks.

We applied Bonferroni correction to determine the *P*-value cutoffs for gene-level association tests. Specifically, when focusing on the common gene set that can be analyzed by all methods, we used the same Bonferroni correction cutoff; otherwise, we used 0.05 divided by the total number of genes tested for each method.

## Results

### Simulation studies

One key advancement of CMO is that it achieves high power to identify new outcome associated genes that could not be identified by competing methods. To evaluate the type I error rate and statistical power under different settings (Methods), we performed simulations studies using randomly selected 10,000 independent White British individuals from UK Biobank (application number 48240). We observed that the type I error rate was well-controlled under different significance levels (Supplementary Table 1). More important, the statistical power of our proposed CMO test was much higher than competing methods when three CpGs and three tissues were associated with the trait (*m* = 3) and CpGs were negatively associated with gene expression (*h*_*c*_ = −1, *h*_*g*_ = 1; Figure 1). The improvement was consistent under different number of causal CpGs and tissues and the effect direction of CpGs (Supplementary Figures 1–4). Interestingly, when CpGs were not associated with the trait directly (i.e., *h*_*c*_ = 0), CMO still achieved comparable power as TWAS and MultiXcan (Supplementary Figures 5 and 6). This is because gene expression may mediate through DNA methylation.

**Figure 1:**
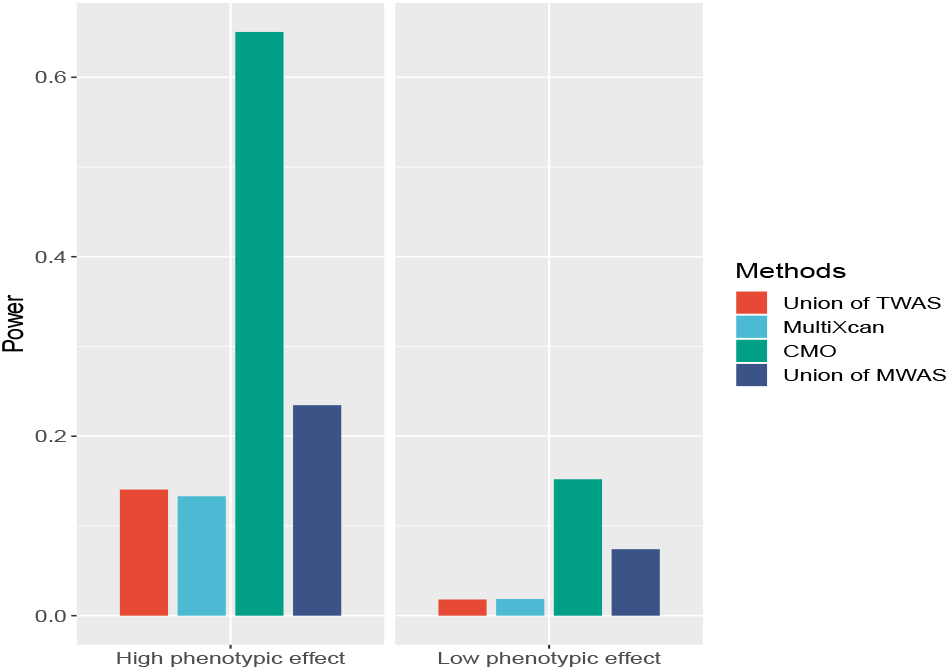
CMO improved statistical power. The left and right panel represent the situations in which genes explain 1% and 0.5% of the total trait variance (heritability, denoted as high/low phenotypic effects). Each method is indicated in the bar plot. We set *m* = 3, *h*_*c*_ = −1, and *h*_*g*_ = 1. The union of TWAS is a method that considers all tissues while applying an additional Bonferroni correction. The union of MWAS is a method that combines results across CpGs in the gene body and promoter regions while applying an additional Bonferroni correction.

### CMO identifies more associations than competing methods

We applied CMO to the summary results from all the 21 GWAS of Susceptibility Weighted Imaging (SWI) reported earlier [38] (*N* = 8, 428) to further evaluate the statistical power of different integrative gene-based tests. The CMO results were compared with those of TWAS [5, 6] and MultiXcan [35]. Figure 3 shows that CMO identified substantially more associations, showing 124.2% improvement compared with TWAS and 91.4% improvement compared with MultiXcan. Additionally, our CMO test identified 268.8% more associated genes than the naive test (Union of MWAS), which combines the results across CpGs in the gene body and promoter regions and applies an additional Bonferroni correction. Different methods test different sets of genes, consequently, we also compared methods over a common set of 11,708 genes that could be analyzed by all the methods. Again, we showed that CMO identified substantially more associations than competing methods (Figure 2). Of note, such improvement was observed consistently across all traits considered (Supplementary Tables 2 and 3).

**Figure 2:**
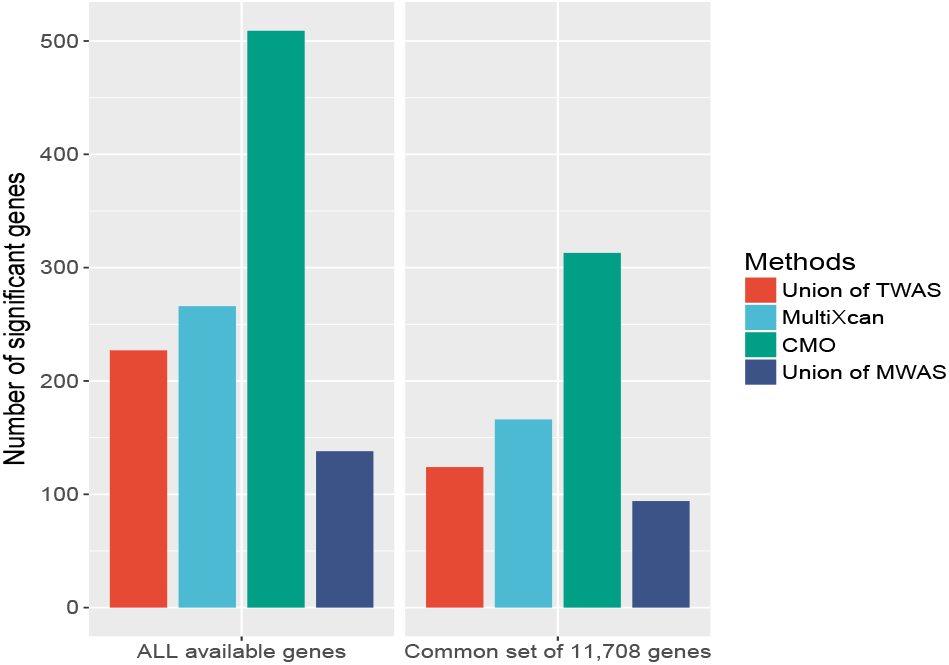
CMO identified more associations for 21 Susceptibility weighted imaging (SWI) related traits. The left and right panel show the numbers of significant genes identified in all available genes and the common set of 11,708 genes, respectively. Each method is indicated in the bar plot. The union of TWAS is a method that considers all tissues while applying an additional Bonferroni correction. The union of MWAS is a method that combines results across CpGs in the gene body and promoter regions while applying an additional Bonferroni correction.

**Figure 3:**
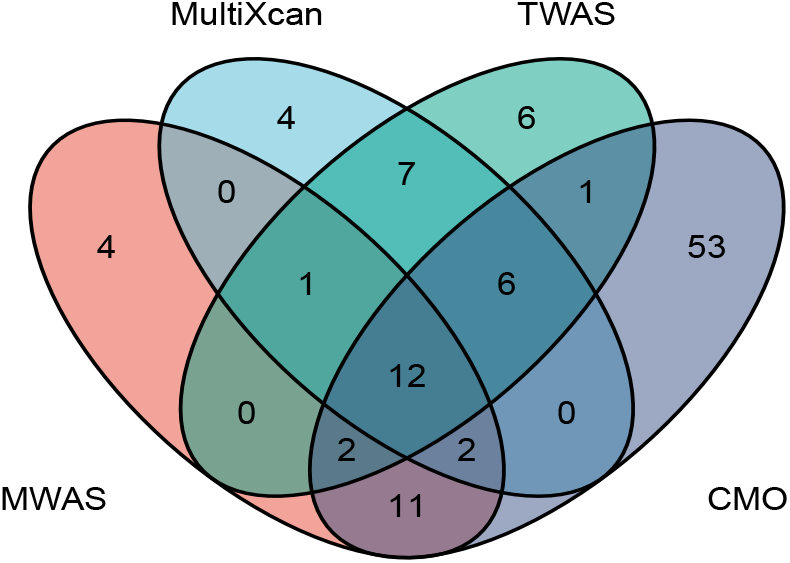
CMO identified more significant genes in the IGAP1 dataset. TWAS stands for TWAS that considers all tissues while applying an additional Bonferroni correction. MWAS is a method that combines results across CpGs in the gene body and promoter regions while applying an additional Bonferroni correction.

### CMO identifies replicable associated genes for AD

We performed a multi-stage gene-level association study for AD (Methods) to further demonstrate CMO’s effectiveness and improve our understanding of the genetic basis of AD. Specifically, we first considered applications to the smaller stage 1 GWAS summary statistics from the International Genomics of Alzheimer’s Project [24] (IGAP1; N = 54,162) for discovery. CMO, Union of MWAS, Union of TWAS, and MultiXcan identified 87, 32, 35, and 32 genome-wide significant genes, respectively (Figure 3). Of note, perhaps because DNA methylation could regulate gene expression and incorporated enhancer-promoter interactions, CMO identified much more significant genes than TWAS and MultiXcan. When we focused on the common set of 11,708 genes that could be analyzed by all methods, CMO still identified substantially more associations than the competing methods (Supplementary Figure 7).

Second, for replication, we used independent summary results from the GWAS by proxy of UK Biobank data [25] (GWAX; *N* = 114, 564). Even though GWAX used the ‘proxy’ AD phenotype based on family history, the replication rate was high: out of 87 significant genes identified by CMO, 12 were replicated under the Bonferroni-corrected significance threshold (*P* = 2.3 × 10^−28^ by a hypergeometric test) and 16 were replicated under a relaxed *p*-value cutoff 0.05 (*P* = 1.2×10^−7^ by a hypergeometric test). Additionally, by searching the GWAS Catalog 1.0 (Version r2019–12–16), we found that 53 out of 87 (60.9%) genes were reported by existing studies (*P* = 6.4 × 10^−50^ by a hypergeometric test). Overall, these two complementary replication efforts demonstrate that the CMO approach is highly robust and particularly more powerful.

### CMO identifies novel associated genes for AD

We next applied CMO to the largest available AD GWAS with up to 455,258 individuals of European ancestry [26] for novel gene discovery. In total, CMO identified 159 genome-wide significant genes (Figure 4 and Supplementary Table 4), which was substantially more than competing methods (Supplementary Figure 8 and Supplementary Tables 5–7). Importantly, 109 out of 159 CMO identified associations were unidentified by MWAS (Supplementary Figure 8), and were driven by the genetically predicted CpGs in enhancer regions. This highlights the importance of integrating enhancer-promoter interaction information in a CMO test.

**Figure 4:**
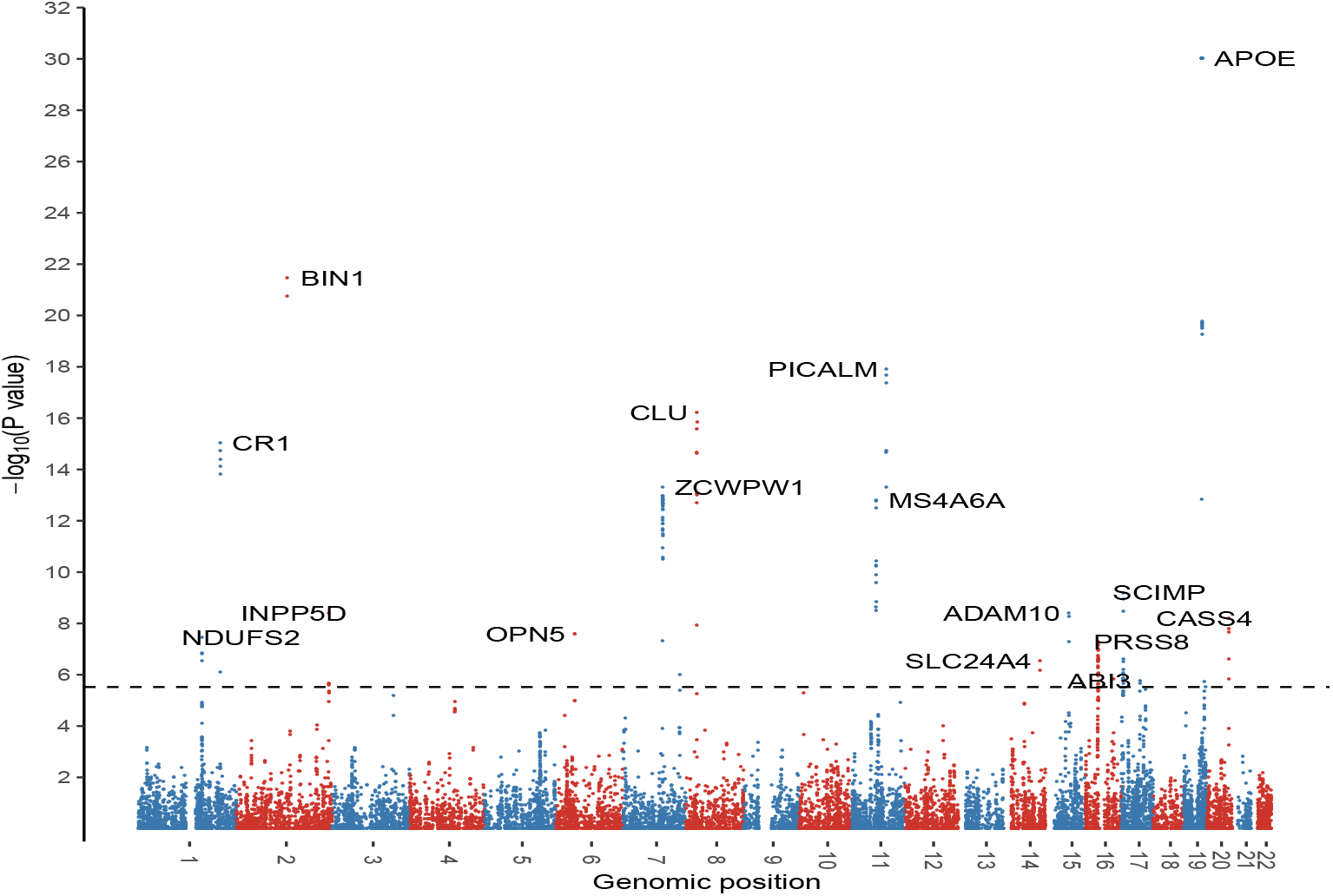
Manhattan plot in a 2019 meta-analyzed AD GWAS dataset [26] (*N* = 455, 258, CMO test). For visualization purposes, *p*-values are truncated at 1 × 10^−30^. The horizontal line marks the genome-wide significance threshold 3 × 10^−6^. The most significant gene at each locus is labeled.

Most significant genes identified by CMO are located at known AD risk loci (Supplementary Table 4) [26, 39]. These include the *CR1* locus in chromosome 1, *BIN1*, *INPP5D* loci in chromosome 2, *CD2AP* locus in chromosome 6, *CLU* locus in chromosome 8, *MS4A6A* locus in chromosome 11, *SLC24A4* locus in chromosome 14, *ADAM10* locus in chromosome 15, *KAT8* locus in chromosome 16, *ABI3* locus inn chromosome 17, *APOE* locus in chromosome 19, and *CASS4* locus in chromosome 20.

Ingenuity pathway analysis (IPA) further validates our findings and provides new insights. For example, the top disease suggested by “Disease and Functions” module in IPA was late-onset AD (*p* = 4.1 × 10^−15^ by a Fisher’s exact test), which involves ten associated genes: *ABI3*, *ADAM10*, *APOE*, *BIN1*, *CD2AP*, *CLU*, *CR1*, *EPHA1*, *MS4A4A*, *PICALM*. The other top suggested diseases were adhesion of blood cells (*p* = 3.4 × 10^−9^) and cancer (*p* = 4.1 × 10^−7^). Cancer and AD have inverse relationship in many common biological mechanism like P53, estrogen, and growth factors [40]. Interestingly, filgrastim, a drug that stimulates the growth of white blood cells, was linked to the identified genes (*p* = 1.5 × 10^−5^. This finding indicates a drug repurposing opportunity for using filgrastim to treat AD, which may deserve further investigation.

Beyond these confirmatory results, we also identified six novel loci for AD, which are at least ±500 kb away from genome-wide significant risk SNPs identified in the 2019 meta-analyzed AD GWAS dataset (including *ZKSCAN5* [*p* = 1.08 × 10^−13^], *FZD3* [*p* = 1.52 × 10^−16^], *BNIP2* [*p* = 5.57 × 10^−9^, *ZNF720* [*p* = 9.22 × 10^−7^, *IL34* [1.60 × 10^−6^], and *ZNF404* [*p* < 1 × 10^−20^]). In comparison, Union of TWAS, MultiXcan, MWAS identified 3, 1, and 0 novel genes, respectively (Supplementary Figure 9). *FZD3* expresses a receptor required for the Wnt signaling pathway [41]. Wnt signaling pathway involves the development of the central nervous system, and the loss of Wnt signaling is associated with the neurotoxicity of the amyloid-*β*. Recall, amyloid-*β* is an established major player for AD pathogenesis [42]. Another example is *IL34*, which is reported to be associated with tau protein, a biomarker for AD [43]. Critically, these six novel genes have not been identified by the competing methods, showcasing the power of our proposed approach.

## Discussion

Integrating genetic regulatory information when performing a gene-level association test usually improves the statistical power because most identified GWAS variants are located in non-coding regions and act by affecting gene regulation. Both simulations and real data applications with summary-level GWAS results demonstrate that the potential power gain of our proposed method CMO. Unlike existing integrative approaches, CMO is the first method to integrate genetically predicted DNAm values and enhancer-promoter interaction with GWAS association results. Compared to TWAS and its extension MultiXcan, our approach generally identifies substantially more associated and overlooked genes for AD, which can be further validated by an independent study. By analyzing data of to date the largest AD GWAS of 71,880 (proxy) cases and 383,378 (proxy) controls of European ancestry, we found associated genes identified by CMO were enriched in lateonset AD pathway and were linked to filgrastim, a drug that stimulates the growth of white blood cells. Furthermore, CMO identified six novel loci for AD, which can be further investigated for improving the risk assessment of AD.

CMO is a computationally efficient method, since *p*-values can be calculated analytically. For example, by a parallel computing strategy with 100 cores in a standard server (each core has 4 GB memory), CMO completed calculating *p*-values for all available genes in IGAP1 data within approximately 1.5 minutes. To facilitate data analysis for both clinical and statistical investigators, we have implemented our proposed method CMO into open-source software.

We view CMO to be complementary (rather than superior) to existing integrative methods. This is because different methods focus on a different set of genes and the underlying genetic structure for each gene is unknown. Thus, each method may be able to identify outcome associated genes that may not be identified by other methods. Specifically, we expect that TWAS will be advantageous when the genetically predicted gene expression is associated with the trait of interest directly without the involvement of DNA methylation influence. On the other hand, when the gene is associated with the trait through DNA methylation pathways, especially those in an enhancer, our proposed method CMO will be more powerful.

We note several limitations of CMO method. First, CMO is an association-based test, and the significant genes identified by CMO do not imply causality. One may apply fine-mapping methods such as FOCUS [44] and FOGS [45] to prioritize putative causal genes. Second, the power of CMO depends on how accurate the genetically predicted DNAm values are. This is similar to the power of TWAS is limited by the quality of imputed gene expression. Currently, the prediction models were provided by [21] and were constructed based on adults of European ancestry. We expect the power of CMO will be further improved when better prediction models are incorporated. Third, for simplicity, all CpGs have the same weight in the current version of CMO. The power may be further improved by incorporating prior information, such as giving more weights for the CpGs in promoter regions. Fourth, we focused on blood tissue for illustration and risk assessment. Similar ideas can be applied to tissue-specific analysis to better understand the etiology of AD. For example, we may use PsychENCODE [46] and ROS/MAP data [47] to construct brain-specific DNAm methylation prediction models and incorporate brain-specific enhancer information. We leave these interesting topics for future research.

In summary, CMO is a novel, robust, and powerful method to perform gene-level association analysis. Enhanced by modern statistical techniques, CMO integrates genetically predicted DNAm values in enhancers, promoters, and the gene body region with GWAS summary results to identify outcome associated genes that may have been ignored by competing methods.

## Declarations

### Availability of data and materials

The data used in this work were obtained from the following publicly available datasets: IGAP1, GWAX, UKBiobank, a 2019 meta-analyzed AD GWAS results, and a imaging-derived phenotype GWAS results. The data resources are summarized in Supplementary Table 8.

We used the publicly available software and tools for competing methods. All codes used to generate results that are reported in this manuscript and software for our newly proposed method CMO are available at https://github.com/ChongWuLab/CMO.

## Acknowledgement

We thank Yanfa Sun for performing IPA analysis. We thank the individuals involved in the UK Biobank and GWAS datasets for their participation and the research teams for their work on collecting, processing, and sharing these datasets. This research has been conducted using the UK Biobank recourse (application number 48240), subject to a data transfer agreement. We acknowledge all of the studies that made their GWAS summary results publicly available.

