## Supplementary Figures and Tables for "A gene-level methylome-wide association analysis identifies novel Alzheimer’s disease genes"

**Department of Statistics**
117 N. Woodward Avenue
P.O. Box 3064330
Tallahassee, Florida 32306-4330
*(850) 644-3218*

June 24, 2020

Julio Licinio, MD, PhD, MBA, MS,
Chief Editor, *Molecular psychiatry*

**RE: A gene-level methylome-wide association analysis identifies novel Alzheimer's disease genes**

Dear Dr. Licinio,

We would like to submit the manuscript referenced above for publication consideration in *Molecular psychiatry*.

In this paper we propose a novel integrative gene-level association test that integrates genome-wide association studies (GWAS) results with emerging/accumulating 1) genetically predicted DNA methylation models and 2) functional regulatory information (i.e., enhancer-promoter inter- actions). This work is motivated by popular transcriptome-wide association studies (TWAS) that leverage transcriptomic reference data to gene-level association tests. By reanalyzing several GWAS summary results, our approach identified an average of 124% more associations than that of conventional TWAS. Importantly, our approach identified candidate genes at six novel Alzheimer's disease loci (including *ZKSCAN5*, *FZD3*, *BNIP2*, *ZNF720*, *IL34*, and *ZNF404*), which can be further investigated for their potential utility in AD risk assessment. These results suggest that our newly proposed approach improves the statistical power. Also, for the first time, we are able to effectively integrate genetically regulated DNA methylation and promoter-enhancer interactions with GWAS results.

We expect there will be significant, broad interest in this paper as novel strategies to improve such -omics integrative analysis are urgently needed and Alzheimer's disease represents an increasing public health burden due to the growing aging population.

In this time of uncertainty related to the COVID-19 pandemic, we realize that the burden of review may be greater than usual. Please know that we greatly appreciate your time consideration of this work, as well as that of the *Molecular psychiatry* reviewers. We sincerely thank you in advance for your time and consideration of our work.

Sincerely,

Chong Wu
Assistant Professor,
Department of Statistics,
Florida State University
